# A novel pH-regulated, unusual 603 bp overlapping protein coding gene *pop* is encoded antisense to *ompA* in *Escherichia coli* O157:H7 (EHEC)

**DOI:** 10.1101/852251

**Authors:** Barbara Zehentner, Zachary Ardern, Michaela Kreitmeier, Siegfried Scherer, Klaus Neuhaus

## Abstract

Antisense transcription is well known in bacteria. However, translation of antisense RNAs is typically not considered, as the implied overlapping coding at a DNA locus is assumed to be highly improbable. Therefore, such overlapping genes are systematically excluded in prokaryotic genome annotation. Here we report an exceptional 603 bp long open reading frame completely embedded in antisense to the gene of the outer membrane protein *ompA*. Ribosomal profiling revealed translation of the mRNA and the protein was detected in Western blots. A σ^70^ promoter, transcription start site, Shine-Dalgarno motif and rho-independent terminator were experimentally validated. A pH-dependent phenotype conferred by the protein was shown in competitive overexpression growth experiments of a translationally arrested mutant *versus* wild type. We designate this novel gene *pop* (**p**H-regulated **o**verlapping **p**rotein-coding gene). Increasing evidence based on ribosome-profiling indicates translation of antisense RNA, suggesting that more overlapping genes of unknown function may exist in bacteria.

## Introduction

Due to the nature of the genetic triplet code, six reading frames exist on the two strands of a DNA molecule. Two genes encoded by two different reading frames at the same DNA locus are called overlapping genes (OLGs). Most overlaps found in bacterial genomes are very short; the majority are only 1 or 4 bp in same-strand orientation (Johnson and Chisholm, 2004) and such overlaps seem to increase fitness (e. g. Saha et al., 2016). In our paper, however, we focus on a non-trivial, extensively overlapping open reading frame (ORF) fully embedded in antisense to an annotated gene. Non-trivially overlapping ORFs in alternative reading frames of the coding strand were discovered in bacteriophage ΦX174 by Barrell *et al*. as early as 1976. Today, the existence of overlapping genes is accepted in viruses, although the driving force for developing gene overlaps is still debated, with theories about size constraint of the genome in the viral capsid, gene novelty, or evolutionary exploration being discussed (Brandes and Linial, 2016, Chirico et al., 2010).

Very little work has been devoted to the exploration of overlapping genes in prokaryotes (Ellis and Brown, 2003, Rogozin et al., 2002). As bacterial genomes are typically much larger than those of viruses, the original hypothesis for the selection pressure associated with the evolution of overlapping genes, namely, increasing the coding capacity in size-restricted genomes (Normark *et al*., 1983), is not valid for prokaryotes. This may be one reason for the lack of knowledge about these amazing gene constructs in bacteria. Furthermore, overlaps of more than 30 amino acids (i.e., non-trivial overlaps) are systematically excluded in genome annotations (Warren et al., 2010). Nevertheless, statistical analysis of bacterial genomes showed that ORFs overlapping annotated genes in alternative reading frames are longer than expected, leading to the hypothesis of a possible selection pressure due to overlapping protein-coding genes (Mir et al., 2012). At least some evidence for functionality of a few non-trivially overlapping genes has been presented (e. g. Balabanov et al., 2012, Behrens et al., 2002). However, the task of functionally characterizing overlapping genes is challenging. It is assumed that these genes originated by overprinting of existing, annotated genes (Sabath et al., 2012) and may constitute an evolutionarily young part of the functional genome of bacteria (Fellner et al., 2014, Fellner et al., 2015). As such, weak expression as well as non-essential function rather than strong phenotypes are expected in general, and sensitive methods are necessary to uncover phenotypic effects caused by the expression of non-trivially overlapping genes.

The human pathogenic bacterium *Escherichia coli* O157:H7 (EHEC) and its genome are well characterized, especially with respect to virulence and the associated diseases like enterocolitis, diarrhea and hemolytic uremic syndrome (Betz et al., 2016, Lim et al., 2010, Stevens and Frankel, 2014). However, the coding capacity of EHEC’s genome is likely to be significantly underestimated, both regarding short intergenic genes (Hücker et al., 2017, Neuhaus et al., 2016) as well as non-trivially overlapping genes (Fellner et al., 2014, Fellner et al., 2015, Hücker et al., 2018a, Hücker et al., 2018b, Vanderhaeghen et al., 2018). Additionally, using a variety of different next generation sequencing based methods (e.g., RNAseq, Cappable-seq, ribosome profiling) evidence has accumulated for widespread antisense transcription and probable translation of these RNAs (Conway et al., 2014, Meydan et al., 2019). If true, this finding would be very significant. Nevertheless, the specificity of the signals found in NGS is sometimes unclear since pervasive translation may occur (Ingolia *et al*., 2014). To gather further evidence for an overlapping coding potential, individual overlapping genes have to be characterized in detail. Such research is in its infancy in bacteria even though, as noted, at least a few overlapping genes have been defined in recent years.

Here, we report on a functional analysis of the unusually long, non-trivially overlapping gene *pop* from *E. coli* O157:H7 strain EDL933, which is completely embedded in antisense to the coding sequence of the conserved outer membrane protein *ompA*.

## Results

### Localization of *pop* in the context of the EHEC genome and its expression

The overlapping gene *pop* from *Escherichia coli* O157:H7 (EHEC) EDL933 probably starts at genome position 1236020 (coordinates following the genome annotation of Latif et al. (2014), GenBank accession CP008957) and has a length of 603 bp (**Fig. 1, Fig. S1**). It is completely embedded in antisense to the coding sequence of the annotated, highly conserved outer membrane protein gene *ompA* (1065 bp), which probably has multiple functions, among which is a porine function and a local cell wall stabilizing action through interaction of OmpA with TolR (Boags et al., 2019). *pop* is located in frame −1 with respect to *ompA* (**Fig. 1 A**). Ribosome profiling of EHEC EDL933 revealed clear evidence of translation of this overlapping gene in LB medium, which is reproducible across biological replicates (**Fig. 2 A, table S1**) and EHEC strains (**Fig. 2 A-C**, see below). Expression of *ompA* is about 150 times higher than *pop*, which is not surprising since OmpA is one of the most highly expressed proteins in *E. coli* (Ortiz-Suarez et al., 2016). The annotated gene *ycbG* (453 bp), encoding a macrodomain ter protein, is located upstream of *pop.* RPKM values of *ycbG* are on average 3 times higher than values of *pop* (**table S1, Fig. 2 D**). The ribosome coverage value (RCV) describes the ‘translational efficacy’ of an RNA molecule 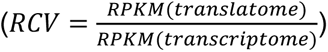. For *pop*, the RCV is reasonably high, greater than 1 in a few instances. According to Neuhaus et al. (2017), transcripts with a RCV higher than 0.355 should be considered to be translated, while untranslated RNAs have a lower RCV. Notably, this measure of the ‘translatability’ of an ORF is on average higher for *pop* than for the annotated upstream gene *ycbG* (**Fig. 2 E, table S1**).

**Figure 1.**
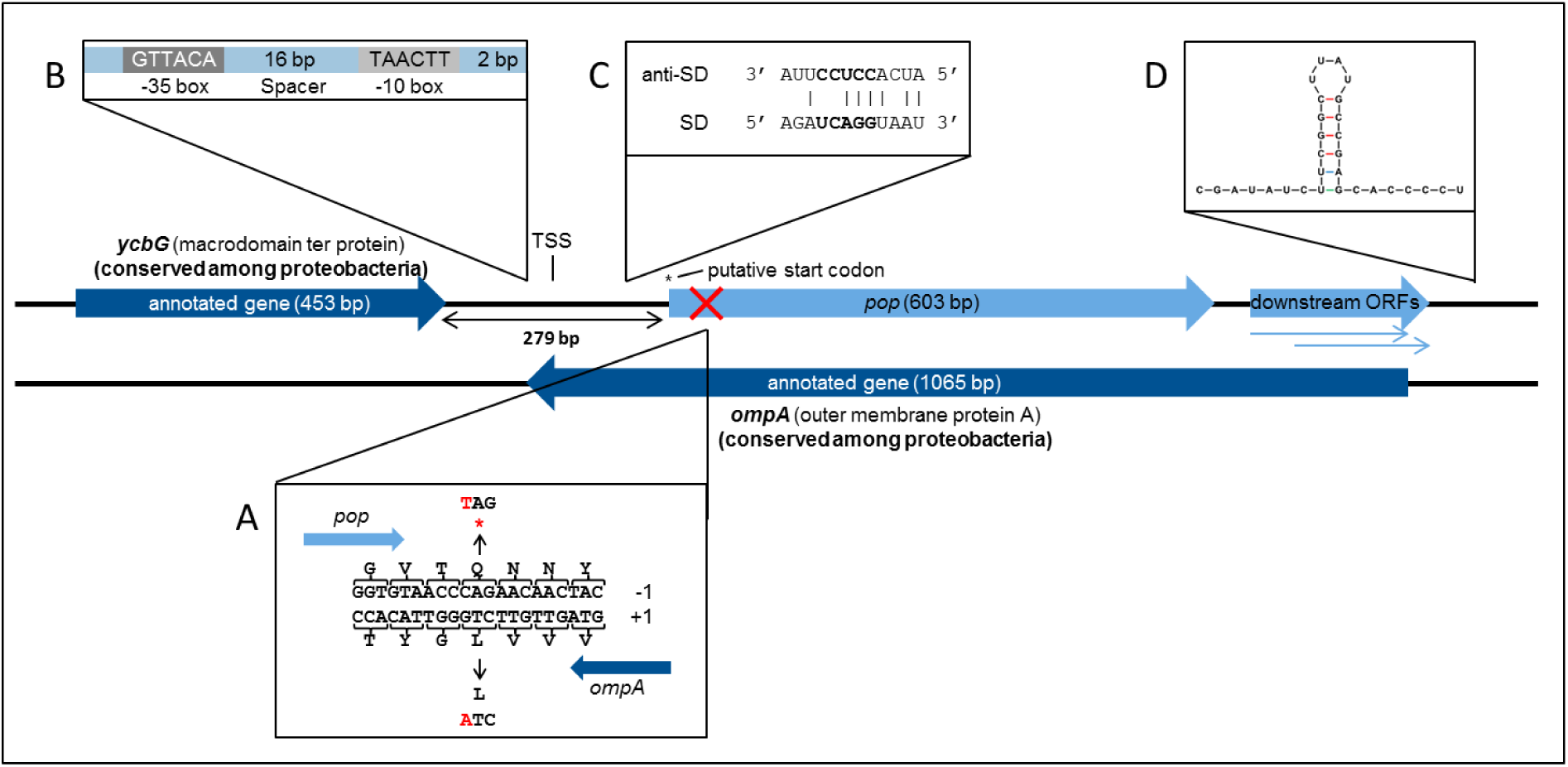
Genomic organization and operon structure of *pop*. *pop* (603 bp) is located downstream of the annotated gene *ycbG* and completely embedded antisense in the sequence of *ompA*. Downstream of *pop* exist two smaller overlapping open reading frames, which overlap *ompA* almost completely and are referred to as downstream ORFs. A transcription start site (TSS) was experimentally identified in the intergenic region of *pop* and *ycbG*. The full genomic sequence of *pop* is given in figure S1. **(A)** Design of a translationally arrested mutant of *pop*. The overlapping ORF *pop* is located in reading frame −1 with respect to *ompA*. Mutants for phenotypic characterization contained a single base substitution C → T at genome position 1236083 as indicated, leading to a stop codon (*) in *pop* and a synonymous change in *ompA*. **(B)** Promoter sequence. The sequences of the −10 box and −35 box as well as the length of the spacer between the conserved boxes and the distance to the TSS are shown. **(C)** Alignment of Shine-Dalgarno (SD) sequence of *pop* and anti-Shine-Dalgarno (anti-SD) sequence. The predicted SD sequence (ΔG° = −3.6 kcal/mol) upstream of the putative start codon is aligned to the consensus of the anti-SD sequence of the 16S rRNA in the 30S ribosomal subunit (Ma *et al*., 2002). The core of the ribosome binding site is displayed in bold letters. **(D)** Secondary structure of the first 40 bp of the predicted terminator. The folding was conducted with Mfold and the structure has a final energy of ΔG = −8.6 kcal/mol.

**Figure 2.**
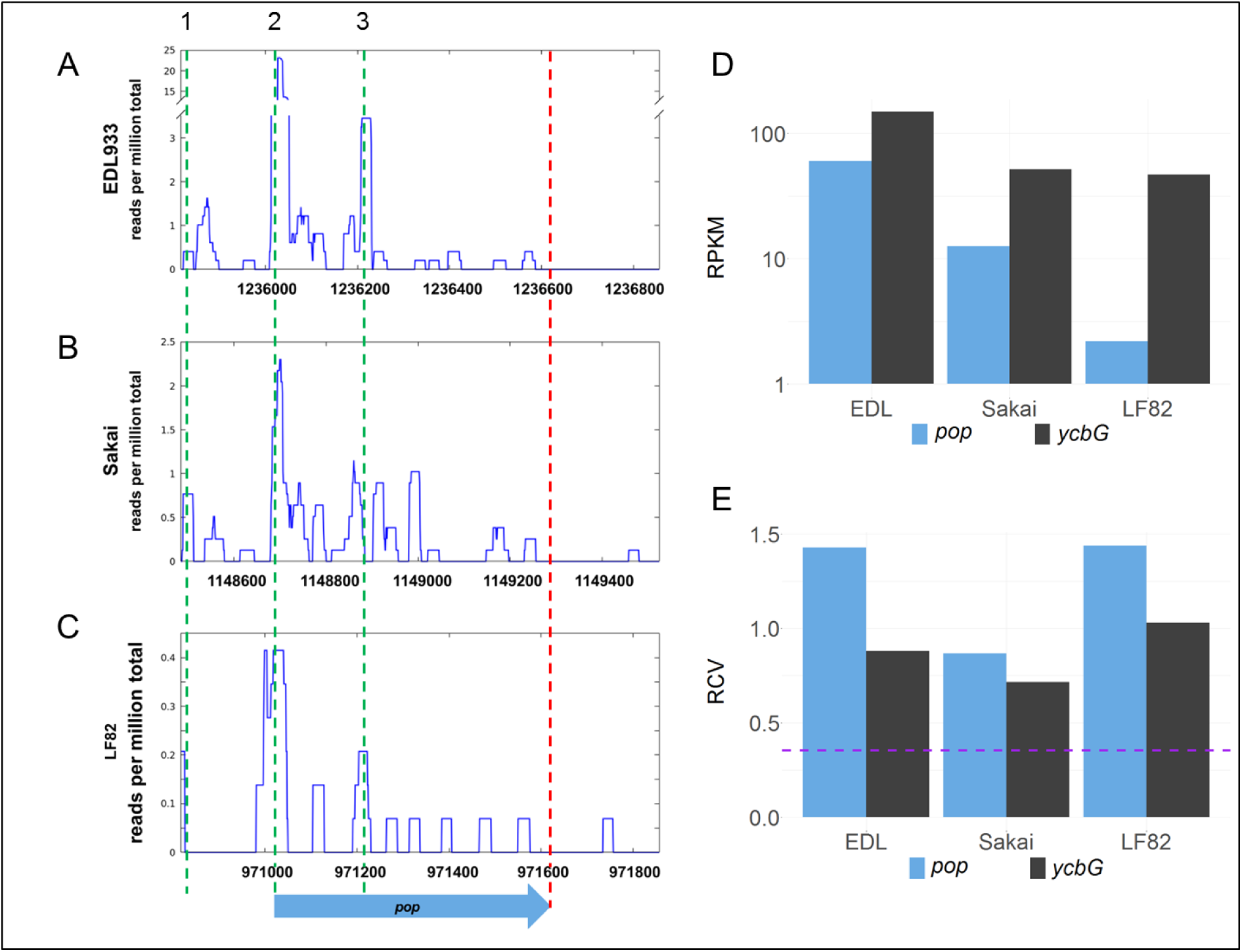
Inter-strain comparison of *pop* translation. Alignment of sequence and ribosomal profiling reads of *pop* and its homologs across pathogenic *E. coli* strains O157:H7 EDL933 **(A)**, O157:H7 Sakai **(B)**, and LF82 **(C)**. Graphs show normalized sequencing reads (RPKM, reads per kilobase per million mapped reads) of ribosome profiling experiments in LB medium **(A, B)** or Schaedler broth **(C)**; the sum signal of two biological replicates is visualized. Three putative start codons are indicated with green dashed lines in region 1 (TTG), 2 (CTG) and 3 (GTG). The stop codon is indicated by a red dashed line. **(D)** Averaged RPKM values of translation and **(E)** ribosomal coverage values (RCV) of overlapping gene *pop* and the upstream annotated gene *ycbG* of three pathogenic strains. Purple dashed line in (E): threshold for translated ORFs, RCV=0.355.

The region between *ycbG* and *pop* contains the transcription start site and a σ^70^ promoter (**Fig. 1 B, details further below**). Two downstream ORFs, which are arranged in frames −1 and −2 with respect to *ompA*, are a little over 200 bp long and mostly overlap with *ompA* (**Fig. 1**). Despite a downstream rho-independent terminator (**Fig. 1 D, details further below**) neither of these ORFs appears to be transcribed or translated (**table S1**) and, therefore, we designate the two ORFs in the following simply as downstream ORFs.

Expression of *pop* was analyzed in three pathogenic *E. coli* strains (O157:H7 EDL933, O157:H7 Sakai, and LF82) and an *E. coli* K12 strain (MG1655; **Fig. 2 A-C, table S1**). Interestingly, *pop* is translated in EDL933, Sakai, and LF82, with highest values in EDL933, whereas it is neither transcribed nor translated in *E. coli* MG1655 indicated by low RPKM values and the RCV. Upstream of the *pop*-ORF, we detected a Shine-Dalgarno sequence (ΔG° = −3.6 kcal/mol) and a possible, but rare start codon CTG nearby, starting at genome position 1236020 (**Fig. 1 C, Fig. S1**).

Additional evidence for the translation initiation site is found in recently published stalled-ribosome profiling data using the antibiotic retapamulin in the strain BL21 (Meydan *et al*., 2019). Five reads are antisense to *ompA*, and all are clustered in the vicinity of the putative start site of *pop* (**Fig. 3 A**). A comparison to weakly expressed annotated genes (selection described in methods section) shows that the putative location of the *pop* translation initiation site is within the typical range for such genes (**Fig. 3 B**), and provides independent evidence locating the start codon within at most a few nucleotides of the predicted site. Similarly, we find that with pooled data from EDL933, using the method of Meydan *et al*. (2019) to predict the ribosomal p-site, described in Methods, precisely identifies the start of the previously mentioned CTG codon (position 1236020) as a translation initiation site.

**Figure 3.**
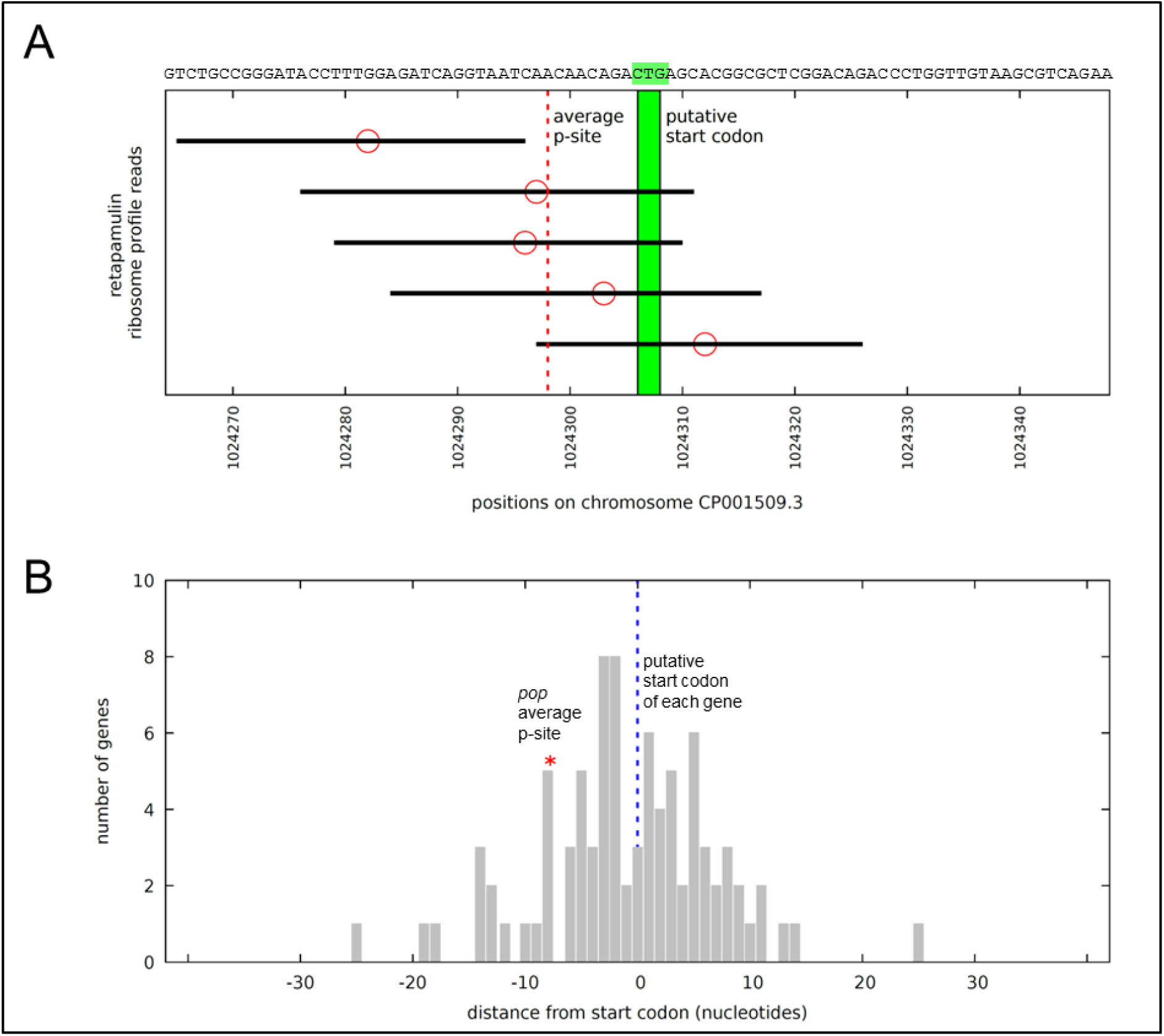
*pop* in stalled-ribosome profiling data of *E. coli* BL21. **(A)** Reads (black bars) antisense to *ompA* in the region of *pop* (sequence indicated above). The five reads situated on the forward strand antisense to *ompA* are shown, aligned to the BL21 chromosome CP001509.3. Ribosomal p-site locations (red circles) are predicted based on the method of Meydan *et al*. (2019) – counting 15 nucleotides from the 3’ end of the read. As there is no clear peak, instead the mean of all of the p-sites was calculated. The mean is shown with a dotted red line, pictured in relation to the putative start codon CTG in green. **(B)** Average ribosomal p-site positions for 85 weakly expressed genes. Positions of average p-sites relative to annotated start codons (blue dotted line), as illustrated in **(A)**, are plotted for all 85 weakly expressed forward (+) strand annotated gene start regions. Weakly expressed is here defined as having at least four mapped reads within 30 nucleotides of the annotated start site, but no single position (peak) with three or more reads. The location of the average p-site for *pop* (red asterisk) lies within this distribution, indicating that the observed cluster of ribosome-stalled reads near the CTG site is informative and that CTG is plausibly the true start codon position.

In summary, *pop* was identified as a translated open reading frame based on ribosome profiling experiments. As its translation signal is reproducible within the *E. coli* strains analyzed, we can reject the hypothesis of stochastic background expression and propose specific expression for *pop*. In the following, we present further data supporting a protein-coding status for the gene as well as expression and functionality of this overlapping gene in the human pathogenic bacterium *E. coli* O157:H7 EDL933.

### Overexpression phenotypes indicate functionality of *pop*

Competitive growth experiments were conducted to analyze the influence of *pop* on EHEC’s growth. For this purpose, *pop* and a translationally arrested mutant ORF were cloned in a low copy overexpression plasmid under the control of an arabinose-inducible promoter (pBAD+*pop* and pBAD+Δ*pop*). The mutant plasmid differs in just one base from the wild type plasmid and this single base substitution introduces a stop codon in the overlapping gene (**Fig. 1 A**) which is neutral in the mother frame of *ompA*. Thus, any difference in growth after overexpression of either the intact or the mutated *pop*-ORF can be explained by the presence or absence of a protein encoded by this overlapping gene.

The competition experiment was conducted in different stress conditions (**Fig. 4 A**). Altered growth of cells overexpressing mutant or wild type sequences was detected in LB-based media supplemented with different stressors, whereas plain LB medium did not have a significant influence on the relative growth of mutant and wild type. For instance, addition of the organic acids L-malic acid and malonic acid as stressors led to better growth of cells containing the wild type plasmid compared to cells expressing the mutated sequence, thus, the presence of *pop* is advantageous in this condition. Addition of the acidic substances resulted in an initial pH shift from 7.4 to 5.8. Reversed proportions were detected when LB was buffered with bicine to a pH of 8.7. However, LB adjusted to acidic (pH = 5.8) or near neutral (pH = 7.4) milieu with the biologic buffers MES and MOPS, respectively, did not result in significant growth differences. Nevertheless, a slightly increased proportion of wild type cells can be seen for MES-buffering, which is in line with expectations based on results in the other acidic conditions. In accordance with the growth advantage of the wild type in the presence of malic acid (**Fig. 4 A**), the mRNA of *pop* was upregulated in the presence of this acid (**Fig. 4 B**).

**Figure 4.**
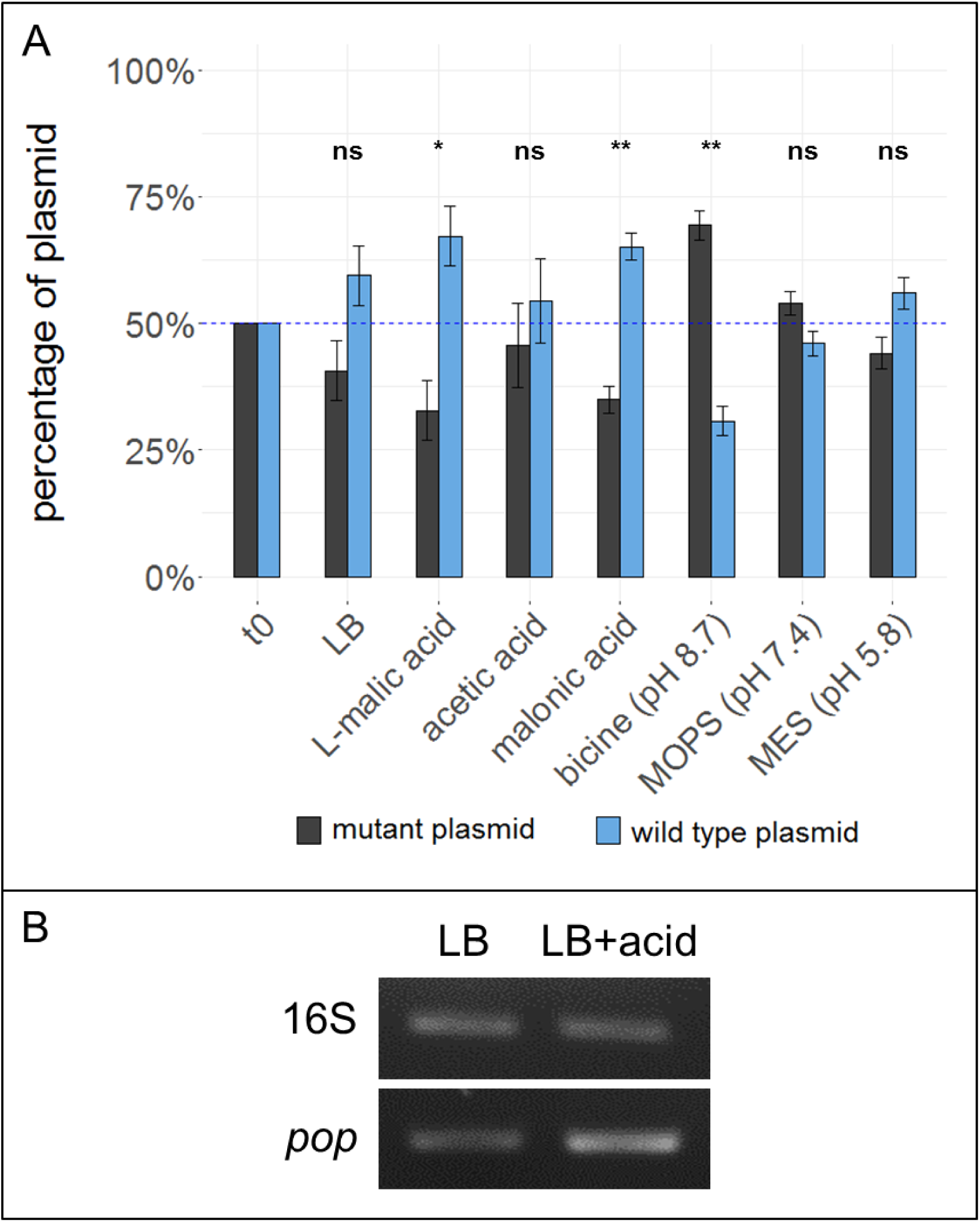
Effect of *pop* expression in various pH ranges. **(A)** Overexpression phenotypes of *pop* in competitive growth assays. Competitive growth of EHEC while overexpressing either intact (pBAD+*pop*) or translationally arrested *pop* (pBAD+Δ*pop*) was conducted in conditions as indicated, i.e. LB medium supplemented with an organic acid or a biological buffer. Mean percentages of wild type (blue bars) or mutated plasmids (grey bars) are shown before (t_0_) and after growth of 22 h. Values are normalized to 50 % input ratio (blue dashed line). Error bars indicate standard deviations. Statistical significance (tested with paired t-test) between wild type plasmids before and after growth is indicated (* p ≤ 0.05; ** p ≤ 0.01; ns, not significant). **(B)** Upregulation of *pop* in the presence of malic acid. Upper lanes: RT-PCR of 16S rRNA, lower lanes: RT-PCR of mRNA of *pop* (primer no. 24+25).

Next, a genomic knock-out for *pop* in *E. coli* O157:H7 EDL933 was constructed. As before, a single base substitution led to a stop codon in *pop*, whereas amino acids in *ompA* remained unchanged (**Fig. 1 A**). We tested the mutant Δ*pop* in several relevant stress conditions in competitive growth against the wild type strain, but did not detect a significant difference in growth in any condition (**Fig. S2**).

Based on the clear effect of overexpression, we propose that *pop* codes for a protein, as mRNAs transcribed from the intact sequence and the translationally arrested variant differ in one nucleotide only. Opposite overexpression phenotypes were found in alkaline buffered and acidified media, so we propose a pH-dependent function. In line with this hypothesis, the mRNA was increased in acidified conditions (**Fig. 4 B**).

### The transcriptional unit of *pop* includes an active promoter and a rho-independent terminator

Cappable-seq (Ettwiller et al., 2016) is a recently developed approach detecting the transcription start site (TSS) of mRNA with next generation sequencing. Using this method, a weak but significant transcriptional start site was determined at genome position 1235862 in the intergenic region between *ycbG* and *pop* in independent biological experiments (**Fig. 1, TSS, Fig. S1**). Two independent bioinformatics tools, BPROM and bTSSfinder, were used to analyze the upstream region of the TSS for potential promoter sequences. Both programs identified a σ^70^ promoter (BPROM LDF score 0.59, Solovyev and Salamov (2011), bTSSfinder score 1.86, Shahmuradov et al. (2017), **Fig. 1 B, Fig. S1**). Although the distance between the transcriptional start site and −10 box of the promoter is not optimal (2 bp instead of approx. 7 bp), promoter sequence activity was verified by means of a GFP-assay (**Fig. 5 A**). We found a significantly enhanced fluorescence in cells harboring the plasmid containing the putative promoter sequence compared to those with the empty vector in LB and bicine-buffered medium. The fluorescence signal of the promoter in the basic milieu (pH 8.7) was strikingly higher, but this may result from GFP accumulation during longer incubation times necessary in this medium (Miller et al., 2000).

**Figure 5.**
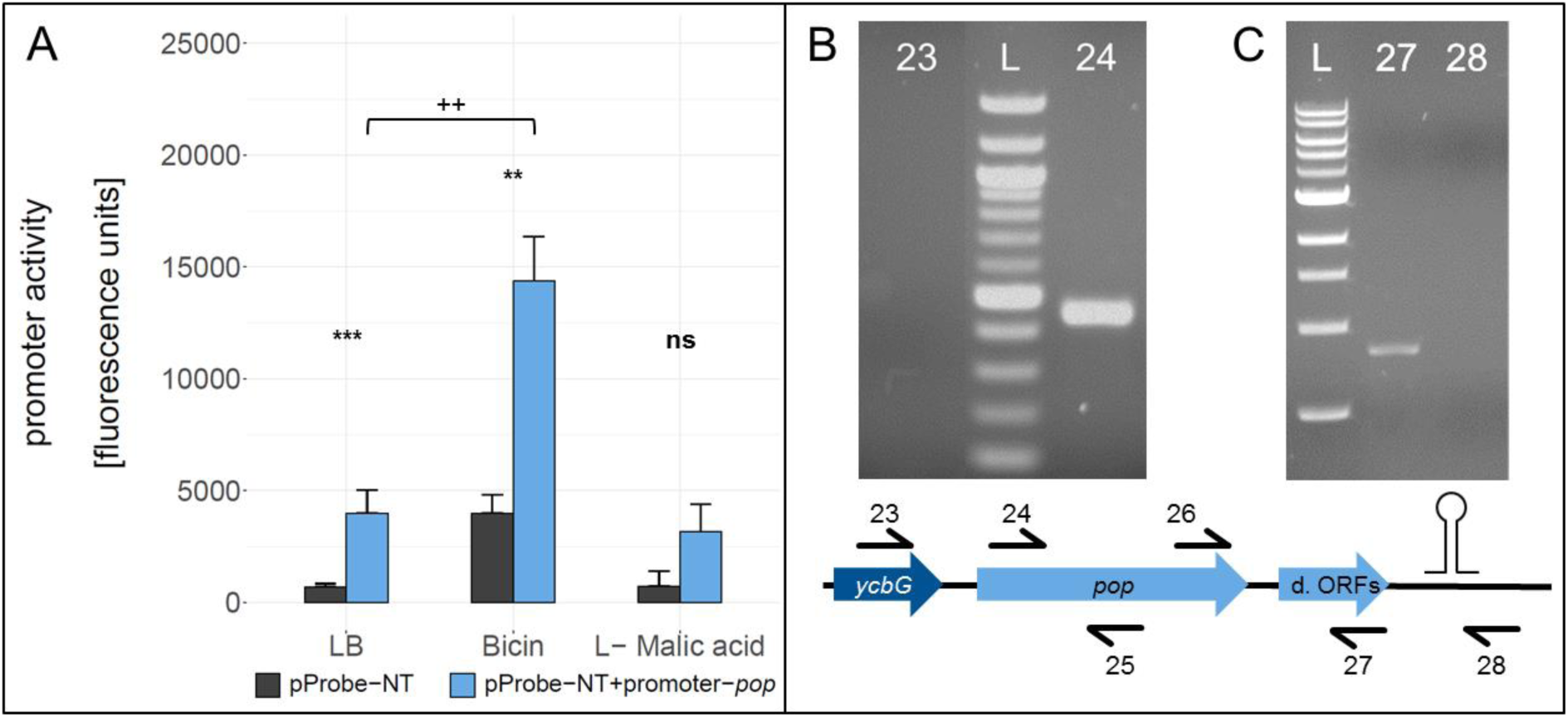
Analysis of the *pop* transcriptional unit. **(A)** Promoter activity assay for the promoter of *pop*, which was introduced in the promoterless GFP vector pProbe-NT. Mean fluorescence units of *E. coli* Top10 cells with the different constructs in culture conditions as indicated are given. Error bars show standard deviations. Statistical significance between empty vector (grey bars) and the promoter construct (blue bars) and between growth conditions was tested with a Welch two sample t-test (**/++ p≤0.01; *** p≤0.001; ns, not significant). **(B)** Test for mono- or polycistronic mRNA. An agarose gel of RT-PCRs is shown. Two different forward primers, binding within *ycbG* (no. 23) or within *pop* (no. 24), were combined with a *pop* reverse primer (no. 25). L: 100 bp DNA Ladder (NEB); 23: PCR with primers 23+25; 24: PCR with primers 24+25. **(C)** Test for the predicted rho-independent terminator. An agarose gel of RT-PCRs is shown. Two different reverse primers, binding upstream (no. 27) or downstream (no. 28) of the stem loop structure, were combined with a *pop* forward primer (no. 26). L, 1 kb DNA Ladder (NEB); 27, PCR with primers 27+26; 28, PCR with primers 28+26; d. ORFs, downstream ORFs.

Since the promoter activity for *pop* is weak compared to promoters of annotated genes, we tested for polycistronic expression starting from the promoter of *ycbG*. Reverse transcription PCR (RT-PCR) was performed to examine the transcript of *pop* (**Fig. 5 B**). No mRNA spanning both genes was detectable, thus, we propose that *pop* is transcribed from the tested promoter monocistronically.

A 120 bp long rho-independent terminator was predicted 295 bp downstream of the stop codon of *pop* using FindTerm (Solovyev and Salamov, 2011). Hypothetical secondary structures of 30-bp segments of this region were created with the tool Quickfold of Mfold (Zuker, 2003). A stable stem loop structure (ΔG = −8.6 kcal/mol) within bases 35 to 78 of the predicted terminator sequence was detected (**Fig. 1 D**). To verify the 3’-end of the mRNA downstream of the hairpin structure, RT-PCRs were performed. We used reverse primers binding either within the downstream ORFs or further downstream, beyond the secondary structure. We observed that *pop* and the downstream ORFs are co-transcribed and transcription is terminated just downstream of the predicted stem loop structure (**Fig. 5 C**).

Based on these results, we conclude that *pop* forms an approximately 1120 bp long transcriptional unit covering almost the entire open reading frame of the mother gene *ompA*, excluding the upstream gene *ycbG* but including the downstream ORFs, ending with a rho-independent terminator.

### Western blot of Pop

Since we detected an active promoter (**Fig. 5 A**) and phenotypes in competitive overexpression experiments (**Fig. 4**), the coding capacity of *pop* was assessed using Western blotting. *pop* was cloned in-frame with an SPA tag (7.7 kDa; following Zuker, 2003) on a pBAD-based plasmid, and overexpressed in EHEC. SPA-tagged proteins were visualized after separating whole cell lysates on tricine gels. The experiment was performed in two different culture conditions (LB at pH 7.4 and bicine-buffered LB at pH 8.7; **Fig. 6**). Besides the expected full-length protein (theoretically 30 kDa, detected approx. 34 kDa), shorter products were immunostained (approx. 20 kDa and 24 kDa). The amount of the full Pop-protein increases within the first 1.5 h after induction and decreases afterwards when overexpressed in LB (**Fig. 6 A**), pointing to an instability of the protein in this condition. In contrast, in bicine-buffered medium, protein expression is somewhat higher at the beginning and does not appear to decrease with time (**Fig. 6 B**). A more detailed evaluation of the smaller products revealed a constant increase in band intensities over time in LB medium, in contrast to a constant signal in bicine. Summarizing, differential *pop* expression is detected by Western blots when growth in LB and bicine is compared.

**Figure 6.**
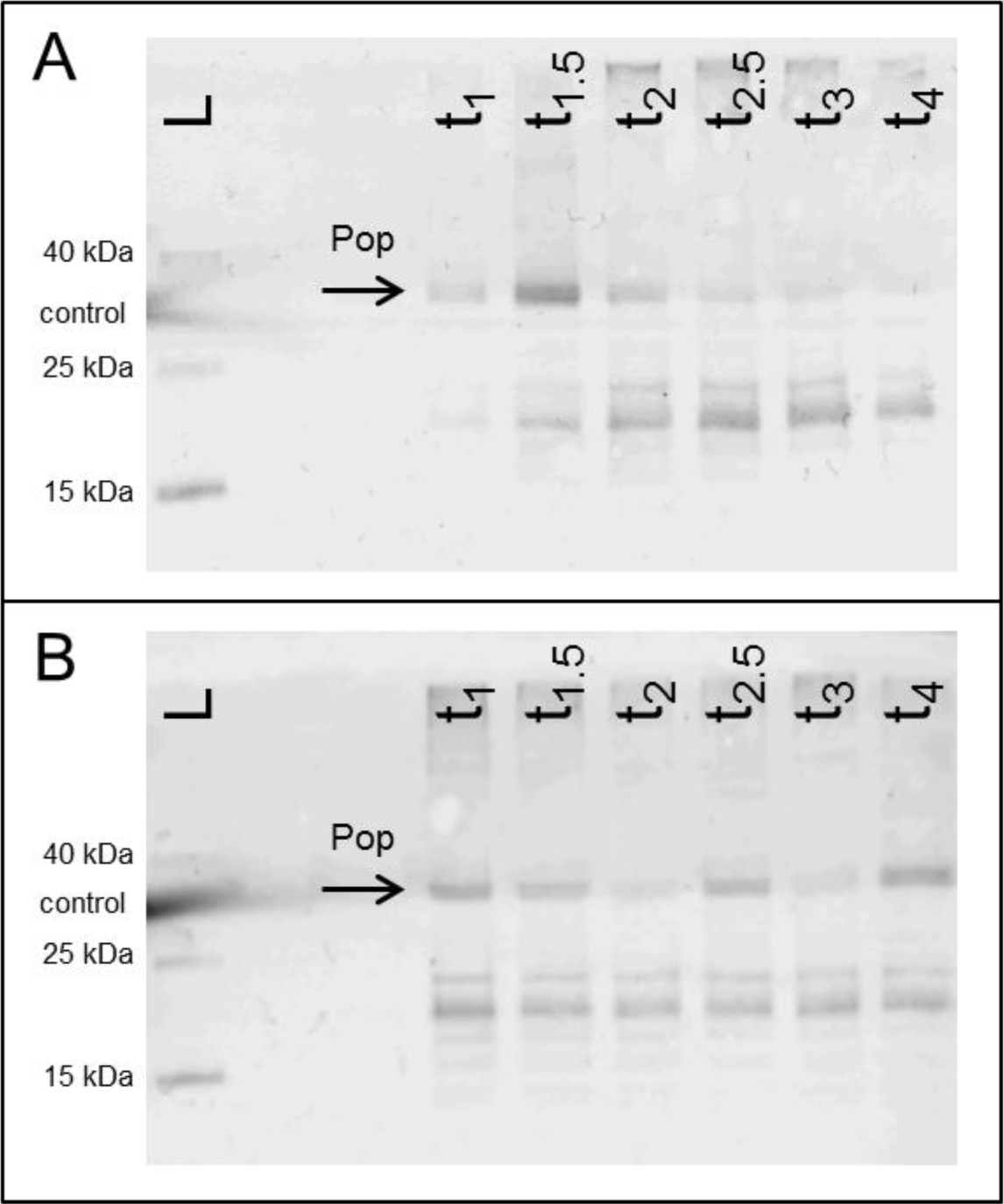
Western blots of Pop protein. Expression of *pop* in frame with a C-terminal SPA-tag. Cells were harvested 1 to 4 h after induction (t_x_) and normalized cell numbers were separated on 16 % tricine gels. The arrow indicates the band of the putative full-length protein Pop. (**A**) The experiment was conducted in LB at pH 7.4 and **(B)** in bicine-buffered LB at pH 8.7. L, Spectra Multicolor Low Range Protein Ladder (Thermo Scientific) with additional internal Western blot control glutathione-S-transferase (band sizes are indicated); t_x_, whole cell extract from samples harvested after x = 1, 1.5, 2, 2.5, 3, and 4 h after induction with arabinose.

### Bioinformatic evidence for *pop* being a protein-coding gene

Protein databases were searched for Pop homologs in order to find hints of a specific function. No significant similarities with annotated proteins were found using blastp analysis in PDB (Protein Data Bank), UniProtKB/Swiss-Prot and the Ref-Seq protein database, but homologous proteins were detected in NCBI’s non-redundant protein sequence (nr) database. However, the hits covered at best 67 % of the amino acid sequence of *pop*. A deeper analysis of the top hit (uncharacterized protein, 67 % coverage, 99 % identity, e-value of 4^−91^) and the genomic sequence of the target organism *Shigella sonnei* showed that its *ompA* homolog was not annotated due to ambiguous bases at its 5’ end, which resulted in a missing start codon for *ompA* in this case. Consequently, *pop* was ‘allowed’ to be predicted *ab initio* as *ompA* had no obvious gene structure and was, therefore, rejected during annotation. This result corroborates the known function of many algorithms like Glimmer, Prodigal or Prokka, which systematically avoid annotation of long overlapping genes (Delcher et al., 2007, Hyatt et al., 2010, Seemann, 2014). Further, NCBI explicitly forbids long overlaps in their prokaryote genome annotation standards (NCBI, 2018). To check whether *pop* is recognized by gene finding algorithms in the case of absent *ompA*, we applied Prodigal to four genomes of bacteria in the family *Enterobacteriaceae* (*Escherichia coli* O157:H7 EDL933, *Shigella dysenteriae*, *Klebsiella pneumoniae*, *Enterobacter cloacae*). Potential start codons of *ompA* were masked with N bases in each genome and consequently *ompA* was not detected. In contrast, *pop* was predicted as a protein-coding gene in all four genomes (**table S2**). The absolute prediction scores of all annotated protein-coding genes in this analysis ranged from −0.5 to > 1000 in EHEC. The total score of *pop* is 14.37 and falls within the lowest 10 % of the 5351 predicted EHEC coding sequences. Nevertheless, sequences with even lower scores than *pop* represent conserved annotated genes, e.g. a fimbrial chaperon or the entericidin A protein, to name only two of many. Thus, *pop* has elements of a gene structure which enable its identification as a protein-coding gene when *ompA* is masked. In the normal case, *pop* is apparently rejected in annotation solely due to its overlapping gene partner *ompA*.

In addition, a homolog for Pop is mentioned in a patent covering the diagnosis and therapy of *Klebsiella pneumoniae* (patent US 6610836, Breton and Osborne, 2003). The homologous protein was found with blastp and has a highly similar amino acid sequence with 99 % coverage and 67 % identity at an e-value of 5×10^−70^. The authors state that all sequences, either nucleic acid or amino acid, including the homolog of Pop, may serve as antigens to diagnose *Klebsiella* infections and allow the production of antibodies against *Klebsiella*. Thus, the patent suggests a function of the Pop homolog in pathogenicity. Interestingly, also in *Klebsiella*, *pop* is fully embedded in its *ompA* mother gene.

## Discussion

Antisense transcription is a widespread phenomenon in bacteria and often connected to regulatory function of the RNAs (Dornenburg et al., 2010, Ettwiller et al., 2016). However, there is increasing evidence that antisense RNAs can be templates for ribosomes to synthesize proteins (Miranda-CasoLuengo et al., 2016, Weaver et al., 2019). So far, characterized non-trivially overlapping genes are typically short (e. g. Fellner et al., 2014, Haycocks and Grainger, 2016, Hücker et al., 2018a); therefore, the discovery and analysis of *pop* with a length of 200 amino acids is of special interest.

The number of coding sequences in bacteria predicted by genome annotation algorithms is underestimated, in particular because neither small genes nor genes with extensive overlap are considered to be true genes (Burge and Karlin, 1998, Delcher et al., 2007, Hücker et al., 2017). Therefore, it is not surprising that *pop* has until now escaped attention. In our study, we detected translation of *pop* in three pathogenic *E. coli* strains, but not in the well-studied *E. coli* K12. The latter finding, in combination with discarding *pop* in automated annotation as it is embedded antisense in the conserved outer membrane protein *ompA*, leads us to presume that it was simply overlooked so far. In contrast, a patent describing polypeptides of *Klebsiella pneumoniae* used for diagnosis explicitly included alternative transcriptional and translational events (Breton and Osborne, 2003) and an expressed homolog of *pop* was found in *Klebsiella,* supporting functionality of *pop* as a protein, possibly having a function in pathogenicity or host response. The latter finding would fit well with the observation of *pop* expression in the pathogenic, but not in the nonpathogenic *E. coli* strain. However, this issue needs further investigation in future studies.

We studied the transcriptional unit of *pop* and identified (i) a TSS (ii) downstream of a σ^70^ promoter, (iii) a potentially coding ORF (i.e., *pop*), and (iv) an experimentally verified rho-independent terminator, but there is uncertainty about the correct start codon of *pop*.

In the ribosome profiling data, we identified three peak regions; a putative start codon of *pop* could be contained in each of these (**regions 1-3 in Fig. 2 A, Fig. S1**). All regions are covered with a substantial number of ribosomal profiling reads, and region 2 is covered best, particularly in EHEC EDL933. We propose that translation for *pop* starts in region 2, especially since a Shine-Dalgarno motif for ribosome binding was predicted and ribosome profiling data across divergent strains point to a putative translation initiation site therein. As mentioned, a nearby CTG is found downstream of the ribosome binding site, representing a rare but sometimes used start codon for prokaryotes (Yamamoto et al., 2018, Hecht et al., 2017, Sussman et al., 1996).

Furthermore, a TTG start codon is present in region 1, representing the longest potential ORF for *pop*. However, we could not find evidence for a TSS or SD-sequence, though the latter is not obligatory for gene expression (Gualerzi and Pon, 2015, Moll et al., 2002). Nevertheless, this TTG was the start codon predicted by Prodigal as the most probable one. Although it has no transcription start site and bicistronic expression along with the upstream gene was excluded in the tested condition, studies showed that read-through of a terminator is common in bacteria in stress conditions (reviewed in Santangelo and Artsimovitch, 2011). Therefore, this TTG start codon could serve as a stress-induced initiation site, which depends on read-through of the transcription termination site of *ycbG*, thus making a TSS unnecessary. To test this hypothesis, future studies using Term-Seq might unveil a transcriptional regulation at the RNA level (Dar *et al*., 2016).

The start codon in region 3 (GTG) is located 45 amino acids downstream of the mutation introduced in *pop* for analysis in competitive growth. This position also cannot be ruled out as a translation initiation site. The growth phenotypes found in competitive growth are not conferred by the protein translated from this start codon, but it might form a protein isoform.

In addition to the gene structure, the Pop protein was analyzed in our study. Western blots verified the protein in two different growth conditions. A condition specific protein stability was assumed as Pop seems to be degraded in LB, while it is stable in bicine buffered medium. Furthermore, the data suggest stable protein isoforms of Pop in bicine, a phenomenon reported for some bacterial proteins recently (Meydan et al., 2019, Vanderhaeghen et al., 2018, Nakahigashi et al., 2016, Waters et al., 2011). Examples are VirF, an AraC like activator in *Shigella*, which possesses two transcriptional start sites and, therefore, two mRNA molecules are templates for translation of the two protein isoforms (Di Martino et al., 2016) and the cobalamin-binding photoreceptor AerR from *Rhodobacter capsulatus* which exists as a long and short protein isoform (Yamamoto *et al*., 2018). Incidentally, translation of the short AerR protein also initiates at a CTG codon. The possibility that Pop isoforms are translated from the same mRNA cannot be excluded, as no internal TSS signal was detected in Cappable-seq. A possible start codon for translation of a smaller isoform could be GTG in region 3, as mentioned in the paragraph before.

Most importantly, competitive overexpression growth assays conducted in this study are the best indication for a proteinaceous nature of the *pop* gene product. As recently shown, not only loss-of-function screenings but also overexpression phenotyping is an appropriate approach to find novel genes and to elucidate their function (Mutalik *et al*., 2019). However, as shown previously, overexpression of unnecessary but usually non-toxic proteins often leads to decreased growth rates (Dong et al., 1995, Shachrai et al., 2010). This could be assumed in our assay conducted in bicine-buffered LB, in which cells expressing the full-length protein had significantly lower growth. Nevertheless, in acidified medium the cells had a growth advantage in comparison to cells expressing the truncated form and, thus, *pop* overexpression is beneficial to EHEC at low pH. This is important since this effect cannot be explained by stressed cells due to protein overexpression. In contrast, analysis of a genomic knock-out suggests that the absence of the protein is not deleterious for EHEC under the conditions tested. While it has been shown that effects of overexpression and knock-out can be complementary, this is not always the case (Prelich, 2012). Several examples exist in which genes can be compensated by each other (e.g. CLN1 and CLN2 in *S. cerevisiae*, Hadwiger et al. (1989); cold shock proteins in bacteria, Xia et al. (2001)). For CLN1 and CLN2, both have similar effects when overexpressed separately, but absence of one of the genes can be balanced out by the other and only a double knock-out has a phenotype.

In summary, we suggest that the investigated open reading frame encodes a protein, since it has overexpression phenotypes in pH stress. Therefore, we propose the name *pop* (**p**H-regulated **o**verlapping **p**rotein-coding gene) for this novel overlapping gene. It should be noted that the *hemC/F/H/L* genes were previously referred to as *popA/B/C/E* but the overlapping gene *pop* is not associated with any function of these. It could be speculated that the positive effect of overexpressed *pop* in acidic medium correlates with the acid tolerance of EHEC necessary to overcome the acidic barrier in the stomach after ingestion (Nguyen and Sperandio, 2012). If true, *pop* could be a pathogenicity or host-environment related gene of EHEC only activated upon specific stress. This fits with the observation that pathogenic *Klebsiella* expresses a Pop homolog (Breton and Osborne, 2003).

Long ORFs embedded antisense to annotated genes like *pop*, as well as other overlapping ORFs, may form a hitherto greatly underestimated source of proteins. Recently developed methods like dRNA-seq (Sharma *et al*., 2010) and Cappable-seq (Ettwiller *et al*., 2016) identified hundreds of transcription start sites antisense to annotated genes producing antisense transcripts with unknown translation status and function. Modern ribosome profiling techniques, including stalling ribosomes at translation initiation sites, identified several unambiguous start codons for protein coding genes which overlap with annotated genes either in sense or in antisense direction (Meydan et al., 2019, Weaver et al., 2019). We suggest that these ‘abnormal’ transcriptional and translational signals in next generation sequencing analysis should not be neglected but analyzed in more detail as has been conducted for the long overlapping gene *pop*, since many novel functions, especially for pathogenicity of novel hosts or survival in new niches, might be ‘hiding’ in the genome of any bacterium.

## Materials and Methods

### Oligonucleotides, bacterial strains and plasmids

All oligonucleotides, bacterial strains and plasmids used or created in this study are listed in table S3.

### Media, media supplements and culture conditions

All *E. coli* strains were cultivated in LB at 37°C, if not stated otherwise. If necessary, medium was supplemented with additives or stressors (see table S4).

### Cloning techniques

Desired sequences were amplified from genomic DNA of *Escherichia coli* O157:H7 EDL933 in a PCR (Q5 polymerase, NEB, Ipswich, Massachusetts, USA) using different primer pairs. PCR fragments were digested with appropriate restriction enzymes (Thermo Fisher Scientific, Waltham, Massachusetts, USA) and ligated in the multiple cloning sites of application specific vectors with T4 DNA ligase (Thermo Fisher Scientific). Vector constructs were transformed in *E. coli* Top10 cells and plated on LB with required antibiotics. Plasmids were isolated (GenElute Plasmid Miniprep Kit, Sigma Aldrich, St. Louis, Missouri, USA) and sequenced with suitable primers (Eurofins Genomics, Ebersberg, Germany) to verify the sequence.

### Creation of translationally arrested knock-out mutants

The genomic knock-out *E. coli* O157:H7 EDL933 Δ*pop* was produced for subsequent competitive growth experiments. Mutation fragments were amplified with primer pairs 1/4 and 2/3 and used in the subsequent overlap extension PCR with primers 3 and 4. The mutation cassette Δ*pop* was cloned in the plasmid pMRS101 using *Apa*I and *Spe*I (selection with ampicillin). The plasmid pMRS101+Δ*pop* was isolated and sequenced with primer 7 and 8. A restriction digest with *Not*I was conducted to remove the high copy ori. The plasmid was re-ligated to the pir-protein dependent low copy plasmid pKNG101+Δ*pop*, propagated into *E. coli* CC118λpir (selection with streptomycin) and isolated. The conjugation strain *E. coli* SM10λpir was transformed with pKNG101+Δ*pop*. Overnight cultures (500 µl) of *E. coli* SM10λpir + pKNG101+Δ*pop* and *E. coli* O157:H7 EDL933 + pSLTS (selection marker ampicillin, temperature sensitive ori) were mixed and cultivated on LB plates (24 h, 30 °C) for conjugation and integration of the plasmid into the genome of EHEC. Conjugated EHEC cells were transferred on LB/ampicillin/streptomycin plates and selectively cultivated (24 h, 30 °C). Correct insertion of the plasmid was confirmed by a PCR using primers 5 and 7. A double resistant strain was used for loop-out of the mutation plasmid. For this, conjugated EHEC + pSLTS was cultivated in LB at 30 °C at 150 rpm until an optical density of OD_600_ = 0.5 and counter-selected on saccharose agar (modified LB without NaCl supplemented with saccharose) containing 0.02 % arabinose to induce the λ red recombination system on pSLTS. A PCR fragment of a streptomycin sensitive clone produced with primers 5 and 6 was sequenced to verify integration of the desired mutation into the chromosome. *E. coli* O157:H7 EDL933 Δ*pop* was cultivated at 37 °C to clear the cells from the plasmid pSLTS.

### Cloning of pBAD+*pop* and pBAD+Δ*pop* for overexpression phenotyping

For overexpression competitive growth, plasmids pBAD+*pop* and pBAD+Δ*pop* were constructed. For the former construct, primers 9 and 10 were used. The latter construct was created similar to the mutation cassette described in the previous section (primers for the mutation fragments 1/9 and 2/10, primers for the mutation cassette: 9 and 10). Both PCR fragments, either wild type or mutant, were cloned in the *Nco*I and *Pst*I sites of pBAD/myc-HisC and plasmids were sequenced with primers 12 and 13. Each of the plasmids was transformed in wild type *E. coli* O157:H7 EDL933 for subsequent competitive growth assays.

### Competitive growth assays

For competitive growth, EHEC overnight cultures of transformants containing pBAD+*pop* or pBAD+Δ*pop* were diluted to OD_600_ = 1 and mixed in equal amounts. Plasmids were isolated from the bacteria mixture and used as time point zero reference. One hundred microliters of a 1:300 dilution of the initial 1:1 bacteria mixture was used to inoculate 10 ml culture medium with appropriate additives (for working concentration of chemicals see table S4; selection marker ampicillin). Overexpression was induced with L-arabinose (0.02 %) at the two time points t_0_ = 0 h and t_x_ = 6.5 h. Plasmids were isolated after t_x_ = 22 h and sequenced with primer 11. Ratios between wild type and mutants were calculated via the amount of mutated and wild type plasmid. The peak heights of mutated and wild type base at the mutated position were measured and normalized to a 1:1 input ratio of the time point zero reference. The percentage of wild type peak and mutant peak were calculated (%_Wt_ = Wt/(Wt + Mt) and %_Mt_ = Mt/(Wt + Mt) with Wt and Mt the peak heights of wild type and mutant plasmid, respectively). Mean values and standard deviations of at least three biologic replicates were calculated. Significance of a possible growth phenotype was tested with a paired t-test between wild type samples of the time point zero reference and the cultured samples (significant result for p-value ≤ 0.05).

Competitive growth of wild type EHEC and translationally arrested mutant *E. coli* O157:H7 EDL933 Δ*pop* was conducted as described above with some exceptions: no selection marker was used; no protein expression was induced; cells were harvested after t_x_ = 18 h; bacterial ratios were determined by sequencing PCR products amplified from cell lysates with primers 5 and 6 for t_0_ and cultured samples (primer for sequencing: 5).

### Construction of overexpression plasmid and Western blot

The plasmid pBAD/myc-HisC was modified to get the overexpression plasmid pBAD/SPA. For this, primers 14 and 15 were annealed (heating at 90 °C, slow cooling) and completed in a PCR where primers 16 and 17 were added after 5 cycles to amplify the fragment. The PCR product was cloned into pBAD/myc-HisC using *Sal*I and *Hind*III restriction enzymes. This resulted in an excision of the myc-epitope and in-frame insertion of the SPA-tag (Zeghouf *et al*., 2004). The sequence of *pop* was cloned after amplification with primers 9 and 13 in the *Nco*I and *Hind*III sites of pBAD/SPA. The plasmid pBAD/SPA+*pop* was sequenced with primers 11 and 12 and transformed into *E. coli* O157:H7 EDL933.

Overexpression was performed in LB medium and bicine-buffered LB medium. Cells were cultivated and protein production was induced with 0.002 % arabinose when an optical density of OD_600_ = 0.3 was reached. Similar cell numbers were harvested at defined time points after induction. Whole cell lysates were prepared by adding 50 µl SDS sample buffer (2 % SDS, 2 % β-mercaptoethanol, 40 % glycerin, 0.04 % coomassie blue G250, 200 mM tris/HCl; pH 6.8) and heating at 95 °C for 10 minutes. Proteins in 10 µl of the lysates were separated on a 16 % tricine gel prepared according to Schägger (2006), and detected afterwards in a Western blot. For this purpose, proteins were blotted semidry (12 V, 20 min) on a PVDF membrane (PSQ membrane, 0.2 µm, Merck Millipore, Burlington, Massachusetts, USA). After incubating the membrane 5 min in 3 % TCA, it was blocked with nonfat dried milk at 4 °C. After three washing steps (TBS-T), the membrane was incubated in a 1:1000 dilution of ANTI-FLAG^®^ M2-Alkaline Phosphatase antibody (Sigma Aldrich) in TBS-T. SPA tagged proteins were visualized with BCIP/NBT.

### Determination of promoter activity by a GFP assay

The promoter sequence of *pop* was amplified with primers 18 and 19. The product was cloned N-terminally into the promoterless GFP-reporter plasmid pProbe-NT using restriction enzymes *Sal*I and *EcoR*I resulting in pProbe-NT+promoter-*pop*. The promoter activity was measured in *E. coli* Top10. For this, 10 ml LB with the appropriate additive (for working concentration of chemicals see table S4; selection marker kanamycin) was inoculated 1:100 with overnight cultures of *E. coli* Top10, *E. coli* Top10+pProbe-NT, and *E. coli* Top10+pProbe-NT+promoter-*pop* and cultivated up to OD_600_ = 0.6. An appropriate number of cells were harvested, washed once and afterwards resuspended in 1xPBS. Fluorescence of 200 µl cell suspension was measured in four technical replicates (Victor3, Perkin Elmer, excitation 485 nm, emission 535 nm, measuring time 1 s). Self-fluorescence of cells was subtracted. Mean values and standard deviation of three independent biological replicates were calculated. Statistically significant differences in the fluorescence of promoter construct and empty plasmid or between promoter constructs in different growth conditions were determined using the Welch two sample t-test (significant result for p-value ≤ 0.05).

### RNA isolation

RNA was isolated from exponentially grown EHEC cultures (OD_600_ = 0.3) using Trizol Reagent (Thermo Fisher Scientific). Cell pellets were resuspended in 600 µl cooled Trizol and cells were disrupted with bead beating using a FastPrep (3-times at 6.5 ms^−1^, 45 s, rest 5 min on ice between the runs). Cooled Chloroform (120 µl) was added, mixed vigorously and incubated 5 min at room temperature. Phases were separated by centrifugation for 15 min (4 °C, 12000×*g*) and total RNA in the aqueous upper phase was precipitated with isopropanol, NaOAc and glycogen (690 µl, 27 µl, and 1 µl, respectively) at −20 °C for 1 h. RNA was pelleted by centrifugation for 10 min and washed twice with 80 % ethanol. Air-dried RNA was resolved in an appropriate volume of RNase-free H_2_O.

### DNase digestion

DNA in RNA samples was digested with Turbo DNase (Ambion) according to the manufacturer. The reaction was stopped with 15 mM EDTA and heating for 10 min at 75 °C. Digested RNA was precipitated with isopropanol, NaOAc and glycogen (690 µl, 27 µl, and 1 µl, respectively) at −20 °C overnight. After centrifugation (20 min, 12000×*g*), the pellet was washed once with 80 % ethanol. Air-dried RNA was resolved in an appropriate volume of RNase-free H_2_O. Successful DNA depletion was verified with a standard PCR using *Taq*-Polymerase (NEB) and primers 21 and 22 binding to the 16S rDNA.

### RT-PCR

DNA-depleted total RNA (500 ng) was used for cDNA synthesis with SuperScript reverse transcriptase (Invitrogen, Thermo Fisher Scientific) according to the manufacturer. A total amount of 25 ng cDNA was used in a standard PCR using *Taq*-Polymerase (NEB; number of cycles for product amplification: 17-20) with primer pairs indicated.

### Bioinformatic analysis

#### Promoter determination

The programs BPROM (Solovyev and Salamov, 2011) and bTSSfinder (Shahmuradov *et al*., 2017) were used to determine the promoter of *pop*. The input sequence for BPROM was 100 bp long and started 65 bp upstream of the identified TSS. The input for bTSSfinder needed to be longer; it spans 300 bp and starts 197 bp upstream of the TSS. BPROM specifies the promoter strength as linear discriminant function (LDF) and a sequence with LDF = 0.2 indicates a promoter with 80 % accuracy and specificity. bTSSfinder calculates scores based on position weight matrices for different sigma factors and accepts promoters greater than the scoring thresholds (0.06 for σ^70^).

#### Terminator analysis

The program FindTerm (Solovyev and Salamov, 2011) was used to analyze 900 bp downstream of *ompA* for a rho-independent terminator (threshold −3). The identified 120 bp long terminator was split into 30 bp segments and all 91 sequences were folded with Mfold (http://www.bioinfo.rpi.edu/applications/mfold; Zuker, 2003) to identify the stem loop structure.

#### Shine-Dalgarno sequence identification

Presence of a Shine-Dalgarno sequence in the region 30 bp upstream of the start codon was analyzed according to Ma et al. (2002). A minimum of ΔG° = −2.9 kcal/mol is required for detection of a ribosome binding site.

#### Gene prediction

Genome sequences and assembly data of *Escherichia coli* O157:H7 str. EDL933 (Accession number CP008957), *Shigella dysenteriae* str. ATCC 13313 (Accession number CP026774.1), *Klebsiella pneumoniae* subsp*. pneumoniae* str. ATCC 13883 (BioProject PRJNA261239) and *Enterobacter cloacae* subsp*. cloacae* str. ATCC 13047 (Accession number CP001918) were downloaded from NCBI. Gene prediction was performed with Prodigal v2.60 (Hyatt *et al*., 2010) with default settings.

#### Ribosomal profiling analysis

Ribosome profiling data of *E. coli* O157:H7 EDL933 (Neuhaus et al. (2017), samples in LB for two biological replicates, SRR5266618, SRR5266620), *E. coli* O157:H7 Sakai (Hücker et al. (2017), sample in LB, SRR5874484; files for the two separate biological replicates were kindly provided by Sarah Hücker) and *E. coli* MG1655 (Wang et al. (2015), samples in LB for two biological replicates; ERR618775, ERR618771) were downloaded from NCBI. Data for *E. coli* LF82 (GenBank accession: NC_011993.1) was produced in our lab according to the methods of Hücker *et al*. (2017) in Schaedler broth medium (anaerobic cultivation). Data evaluation was conducted as following: Adapters were trimmed with cutadapt (Martin, 2011) with a minimum quality score of 10 (q 10) and minimum length of 12 nucleotides (m 12). The trimmed reads were subsequently aligned to the reference chromosome using bowtie2 (Langmead and Salzberg, 2012) in local alignment mode, with zero mismatches (N 0) and a seed length of 19 (L 19). Reads overlapping ribosomal and tRNAs were removed using bedtools (Quinlan and Hall, 2010). Read counts, RPKMs, and coverage were then calculated with respect to the filtered BAM files, using bedtools and a custom bash script.

Stalled-ribosome profiling data from the *E. coli* strain BL21 was obtained from Meydan *et al*. (2019). The adapter sequence was predicted using DNApi.py (Tsuji and Weng, 2016), and adapter trimming, alignment, and removal of rRNAs and tRNAs was conducted as described above. The positions of all reads mapped to the forward strand were obtained using SAMtools (Li *et al*., 2009) and the “bamtobed” tool from BamTools (Barnett et al., 2011).

Reads with predicted ribosomal p-sites within 30 nucleotides in each direction of an annotated forward-strand gene start codon (“start region”) were extracted. Weakly expressed annotated genes with no single position (peak) represented by three or more reads, and also with at least four reads situated within the start region, were found using a custom bash script.

## Author Contributions

B.Z. performed experimental analysis on *pop* in EHEC EDL933 and database searches. Z.A conducted analysis of ribosomal profiling data. M.K. performed ribosomal profiling in LF82. The study was supervised by S.S. and K.N. The first draft of the ms including the figures was written by B.Z. with the help of K.N. and S.S. The final manuscript was read and approved by all authors.

## Acknowledgements

We thank Romy Wecko for excellent technical assistance and Christopher Huptas for his support in gene prediction. We thank Sarah Hücker for providing ribosomal profiling data for the two biological replicates of *E. coli* O157:H7 Sakai. This work was supported in part by the Deutsche Forschungsgemeinschaft (DFG) to S.S. (SCHE316/3-1,2,3).

---

**Figure S1.**
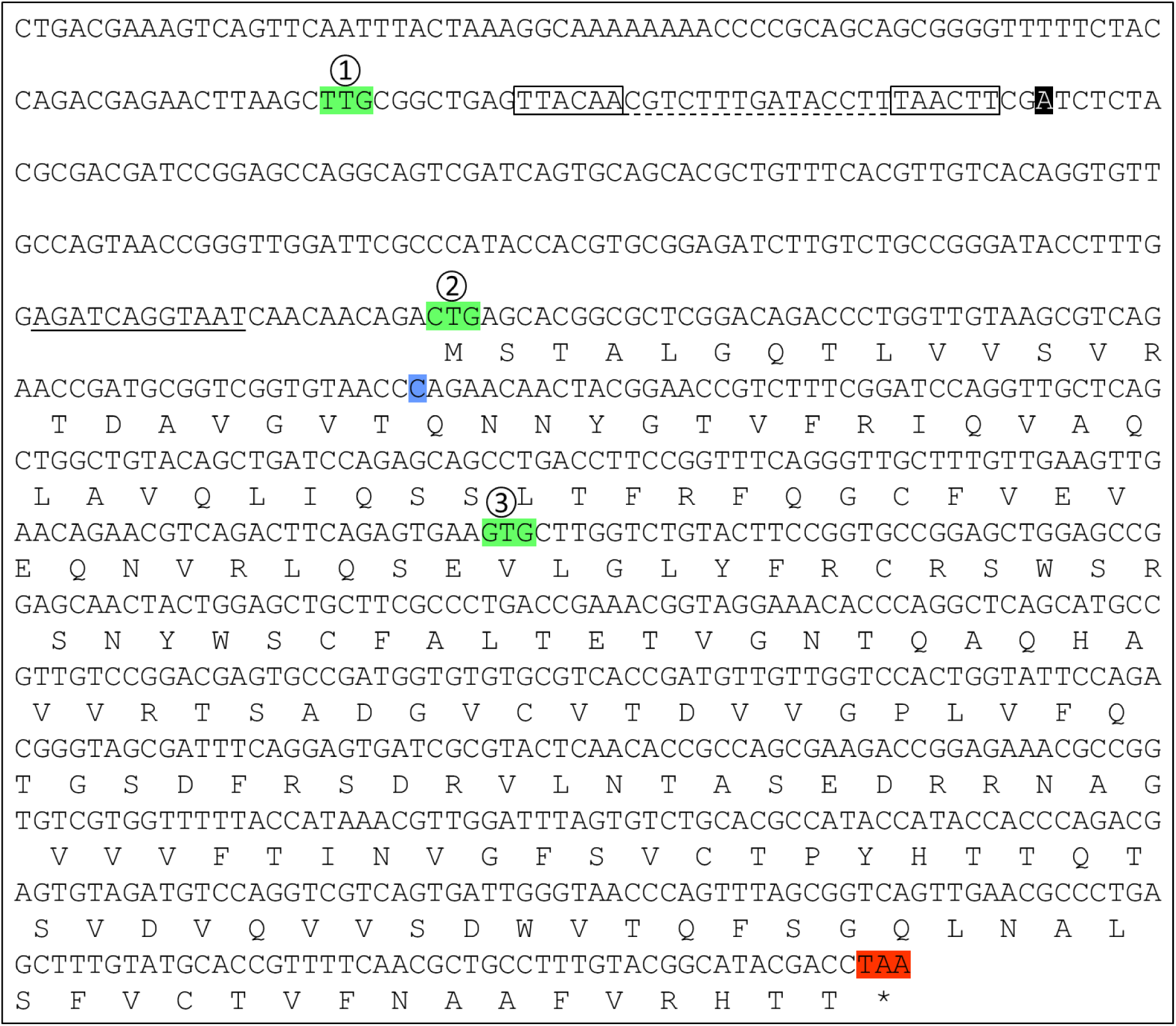
Genomic sequence of *pop*. Nucleic acid and proposed amino acid sequence of *pop* are shown. Several features are highlighted: black, TSS; green, possible NTG start codons (1, 2, 3); red, stop codon (*); blue, mutated position (C → T) resulting in a premature stop codon in *pop*. Black boxes and dashed line indicate the predicted promoter with −35 and −10 box and spacer region, respectively. The Shine-Dalgarno sequence upstream of start codon 2 is underlined.

**Figure S2.**
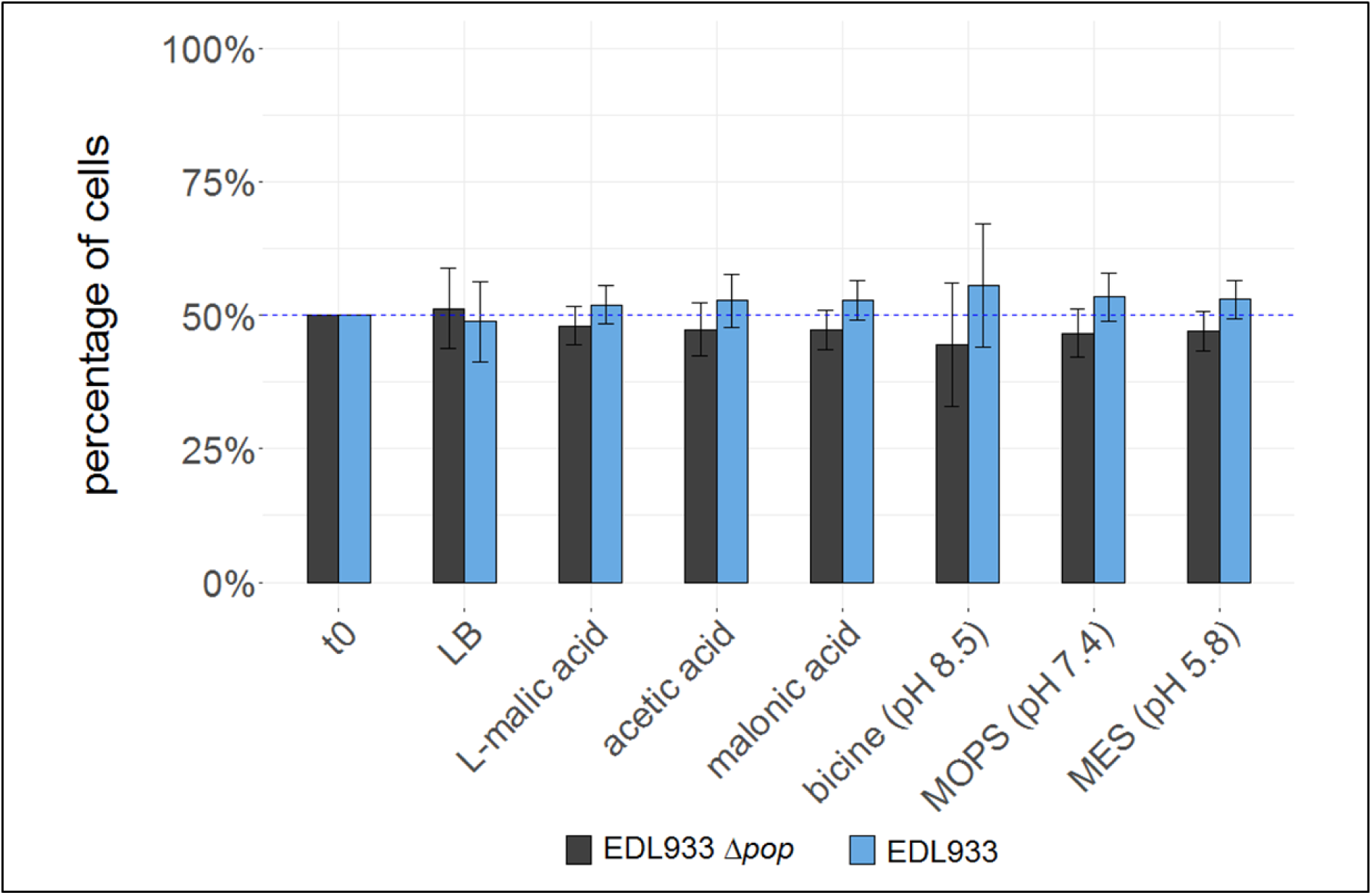
Competitive growth using a genomic knock-out mutant. The genomic knock-out (EDL933 Δ*pop*, grey bars) was grown together with EHEC wild type (EDL933, blue bars) in plain LB or indicated stress conditions (LB medium either supplemented with organic acids or biological buffers). Mean percentages of wild type or mutant cells are shown before (t0) and after growth of 22 h. Values are normalized to 50 % input ratio (blue dashed line). Error bars indicate standard deviations.

**Table S1.**

**Evaluation of ribosome profiling and RNAseq.** Chromosome coordinates (E, F, G and N, O, P) of the open reading frames (H and Q) considered in this study. For each strain (A, B; EDL933, Sakai, LF82, MG1655) in the analyzed ribosomal profiling (C, D) and RNAseq (L, M) datasets, read counts (I, R), RPKM (J, S), and coverage (K, T) are given. Column U, RCV values (i.e. RPKM ribosome profiling / RPKM RNAseq)

**Table S2.**

***pop* prediction with Prodigal.** For each species (A, EHEC EDL933, *Shigella dysenteriae*, *Klebsiella pneumoniae*, *Enterobacter cloacae*) chromosome coordinates (B, C, D) of identified open reading frames are given. The best hit is shaded grey, respectively. Total score (E), coding potential (F), start score (G), start codon (H), RBS motif (I), spacer (J), RBS score (K), upstream score (L), type score (M), and GC concent of the ORF (N) are listed.

**Table S3.**

**Primers, bacterial strains and plasmids used in this study.**

**Table S4.**

**Cultivation stressors, antibiotics and media supplements with corresponding concentrations used in this study.**

